# Adaptive phenotypic plasticity stabilizes evolution in fluctuating environments

**DOI:** 10.1101/2021.05.25.445672

**Authors:** Alexander Lalejini, Austin J. Ferguson, Nkrumah A. Grant, Charles Ofria

**Affiliations:** BEACON Center for the Study of Evolution in Action, Michigan State University, East Lansing, MI, USA; Ecology, Evolution, and Behavior Program, Michigan State University, East Lansing, MI, USA; Department of Computer Science and Engineering, Michigan State University, East Lansing, MI, USA; Department of Biological Sciences, University of Idaho, Moscow, ID, USA

**Keywords:** phenotypic plasticity, experimental evolution, digital evolution, changing environments, Avida

## Abstract

Fluctuating environmental conditions are ubiquitous in natural systems, and populations have evolved various strategies to cope with such fluctuations. The particular mechanisms that evolve profoundly influence subsequent evolutionary dynamics. One such mechanism is phenotypic plasticity, which is the ability of a single genotype to produce alternate phenotypes in an environmentally dependent context. Here, we use digital organisms (self-replicating computer programs) to investigate how adaptive phenotypic plasticity alters evolutionary dynamics and influences evolutionary outcomes in cyclically changing environments. Specifically, we examined the evolutionary histories of both plastic populations and non-plastic populations to ask: (1) Does adaptive plasticity promote or constrain evolutionary change? (2) Are plastic populations better able to evolve and then maintain novel traits? And (3), how does adaptive plasticity affect the potential for maladaptive alleles to accumulate in evolving genomes? We find that populations with adaptive phenotypic plasticity undergo less evolutionary change than non-plastic populations, which must rely on genetic variation from *de novo* mutations to continuously readapt to environmental fluctuations. Indeed, the non-plastic populations undergo more frequent selective sweeps and accumulate many more genetic changes. We find that the repeated selective sweeps in non-plastic populations drive the loss of beneficial traits and accumulation of maladaptive alleles via deleterious hitchhiking, whereas phenotypic plasticity can stabilize populations against environmental fluctuations. This stabilization allows plastic populations to more easily retain novel adaptive traits than their non-plastic counterparts. In general, the evolution of adaptive phenotypic plasticity shifted evolutionary dynamics to be more similar to that of populations evolving in a static environment than to non-plastic populations evolving in an identical fluctuating environment. All natural environments subject populations to some form of change; our findings suggest that the stabilizing effect of phenotypic plasticity plays an important role in subsequent adaptive evolution.

## 1 Introduction

Natural organisms employ a wide range of evolved strategies for coping with environmental change, such as periodic migration (Winger et al., 2019), bet-hedging (Beaumont et al., 2009), adaptive tracking (Barrett and Schluter, 2008), and phenotypic plasticity (Ghalambor et al., 2007). The particular mechanisms that evolve in response to fluctuating environments will also shift the course of subsequent evolution (Wennersten and Forsman, 2012; Schaum and Collins, 2014). As such, if we are to understand or predict evolutionary outcomes, we must be able to identify which mechanisms are most likely to evolve and what constraints and opportunities they impart on subsequent evolution.

In this work, we focus on phenotypic plasticity, which can be defined as the capacity for a single genotype to alter phenotypic expression in response to a change in its environment (West-Eberhard, 2003). Phenotypic plasticity is controlled by genes whose expression is coupled to one or more environmental signals, which may be either biotic or abiotic. For example, the sex ratio of the crustacean *Gammarus duebeni* is modulated by changes in photoperiod and temperature (Dunn et al., 2005), and the reproductive output of some invertebrate species is heightened when infected with parasites to compensate for offspring loss (Chadwick and Little, 2005). In this study, we conducted digital evolution experiments to investigate how the evolution of adaptive phenotypic plasticity shifts the course of evolution in a cyclically changing environment. Specifically, we examined the effects of adaptive plasticity on subsequent genomic and phenotypic change, the capacity to evolve and then maintain novel traits, and the accumulation of deleterious alleles.

Evolutionary biologists have long been interested in how evolutionary change is influenced by phenotypic plasticity because of its role in generating phenotypic variance (Gibert et al., 2019). The effects of phenotypic plasticity on adaptive evolution have been disputed, as few studies have been able to observe both the initial patterns of plasticity and the subsequent divergence of traits in natural populations (Ghalambor et al., 2007; Wund, 2012; Forsman, 2015; Ghalambor et al., 2015; Hendry, 2016). In changing environments, adaptive phenotypic plasticity provides a mechanism for organisms to regulate trait expression within their lifetime, which can stabilize populations through those changes (Gibert et al., 2019). In this context, the stabilizing effect of adaptive plasticity has been hypothesized to constrain the rate of adaptive evolution (Gupta and Lewontin, 1982; Ancel, 2000; Huey et al., 2003; Price et al., 2003; Paenke et al., 2007). That is, directional selection may be weak if environmentally-induced phenotypes are close to the optimum; as such, adaptively plastic populations may evolve slowly (relative to non-plastic populations) unless there is a substantial fitness cost to plasticity.

Phenotypic plasticity allows for the accumulation of genetic variation in genomic regions that are unexpressed under current environmental conditions. Such cryptic (“hidden”) genetic variation can serve as a source of diversity in the population, upon which selection can act when the environment changes (Schlichting, 2008; Levis and Pfennig, 2016). It remains unclear to what extent and under what circumstances this cryptic variation caches adaptive potential or merely accumulates deleterious alleles (Gibson and Dworkin, 2004; Paaby and Rockman, 2014; Zheng et al., 2019).

The “genes as followers” hypothesis (also known as the “plasticity first” hypothesis) predicts that phenotypic plasticity may facilitate adaptive evolutionary change by producing variants with enhanced fitness under stressful or novel conditions (West-Eberhard, 2003; Schwander and Leimar, 2011; Levis and Pfennig, 2016). Environmentally-induced trait changes can be refined through selection over time (*i.e*., genetic accommodation). Further, selection may drive plastic phenotypes to lose their environmental dependence over time in a process known as genetic assimilation (West-Eberhard, 2005; Pigliucci, 2006; Crispo, 2007; Schlichting and Wund, 2014; Levis and Pfennig, 2016). In this way, environmentally-induced phenotypic changes can precede an evolutionary response.

Phenotypic plasticity may also “rescue” populations from extinction under changing environmental conditions by buffering populations against novel stressors. This buffer promotes stability and persistence and grants populations time to further adapt to rapidly changing environmental conditions (West-Eberhard, 2003; Chevin and Lande, 2010).

Disparate predictions about how phenotypic plasticity may shift the course of subsequent evolution are not necessarily mutually exclusive. Genetic and environmental contexts determine if, and to what extent, phenotypic plasticity promotes or constrains subsequent evolution. Figure 1 overviews how we might expect different forms of phenotypic plasticity to result in different evolutionary responses after an environmental change.

**Figure 1.**
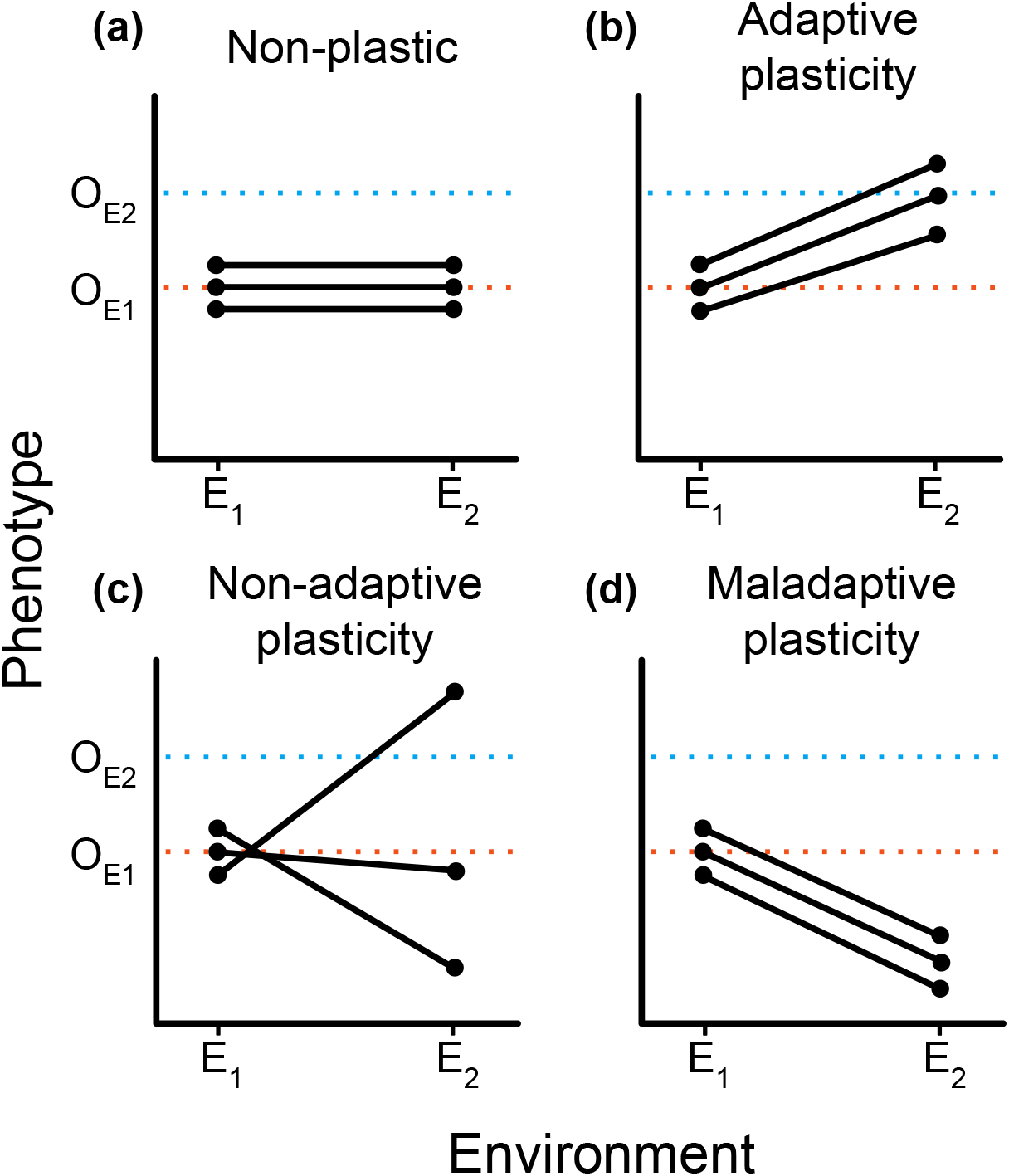
Hypothetical reaction norms for genotypes placed in different environments. In all panels, two environments (denoted E_1_ and E_2_) are shown on the x-axis. The y-axis indicates the phenotype expressed in each environment with O_E1_ and O_E2_ designating the optimal phenotype for E_1_ and E_2_, respectively. Each pair of points connected by a solid black line denotes a genotype, with the points themselves representing its hypothetical phenotypes in each environment. We present four scenarios for how populations could respond to a change from E_1_ to E_2_. (a) A non-plastic population where phenotypes do not change with environmental shifts. In such cases, we would expect strong directional selection toward O_E2_ after the environment changes. (b) An adaptively plastic populations where phenotypes dynamically adjust to the new optimum whenever the environment shifts. As such, we would expect this population to remain relatively stable after the environment changes. (c) A population exhibiting non-adaptive plasticity with substantial variation in how individuals respond to the environmental change. In this case, we expect the change in environment to result in a rapid evolutionary sweep by genotypes closest to the new optimal phenotype. (d) A population exhibiting maladaptive plasticity relative to the given environmental change. When the environment changes, there is little variation for selection to act on, and without beneficial mutations, this population could be at risk of extinction.

Experimental studies investigating the relationship between phenotypic plasticity and evolutionary outcomes can be challenging to conduct in natural systems. Such experiments would require the ability to irreversibly toggle plasticity followed by long periods of evolution during which detailed phenotypic data would need to be collected. Digital evolution experiments have emerged as a powerful research framework from which evolution can be studied. In digital evolution, self-replicating computer programs (digital organisms) compete for resources, mutate, and evolve following Darwinian dynamics (Wilke and Adami, 2002). Digital evolution studies balance the speed and transparency of mathematical and computational simulations with the open-ended realism of laboratory experiments. Modern computers allow us to observe many generations of digital evolution at tractable time scales; thousands of generations can take mere minutes as opposed to months, years, or millennia. Digital evolution systems also allow for perfect, non-invasive data tracking. Such transparency permits the tracking of complete evolutionary histories within an experiment, which circumvents the historical problem of drawing evolutionary inferences using incomplete records (from frozen samples or fossils) and extant genetic sequences. Additionally, digital evolution systems allow for experimental manipulations and analyses that go beyond what is possible in wet-lab experiments. Such analyses have included exhaustive knockouts of every site in a genome to identify the functionality of each (Lenski et al., 2003), comprehensive characterization of local mutational landscapes (Lenski et al., 1999; Canino-Koning et al., 2019), and the real-time reversion of all deleterious mutations as they occur to isolate their long-term effects on evolutionary outcomes (Covert et al., 2013). Furthermore, digital evolution studies allow us to directly toggle the possibility for adaptive plastic responses to evolve, which enables us to empirically test hypotheses that were previously relegated to theoretical analyses.

In this work, we use the Avida Digital Evolution Platform (Ofria et al., 2009). Avida is an open-source system that has been used to conduct a wide range of well-regarded studies on evolutionary dynamics, including the origins of complex features (Lenski et al., 2003), the survival of the flattest effect (Wilke et al., 2001), and the origins of reproductive division of labor (Goldsby et al., 2014). Our experiments build directly on previous studies in Avida that characterized the *de novo* evolution of adaptive phenotypic plasticity (Clune et al., 2007; Lalejini and Ofria, 2016) as well as previous work investigating the evolutionary consequences of fluctuating environments for populations of non-plastic digital organisms (Li and Wilke, 2004; Canino-Koning et al., 2019). Of particular relevance, Clune et al. (2007) and Lalejini and Ofria (2016) experimentally demonstrated that adaptive phenotypic plasticity can evolve given the following four conditions (as identified by Ghalambor et al. 2010): (1) populations experience temporal environmental variation, (2) these environments are differentiable by reliable cues, (3) each environment favors different phenotypic traits, and (4) no single phenotype exhibits high fitness across all environments. We build on this previous work, but we shift our focus from the evolutionary causes of adaptive phenotypic plasticity to investigate its evolutionary consequences in a fluctuating environment.

Each of our experiments are divided into two phases: in phase one, we precondition sets of founder organisms with differing plastic or non-plastic adaptations; in phase two, we examine the subsequent evolution of populations founded with organisms from phase one under specific environmental conditions (Figure 2). First, we examine the evolutionary histories of phase two populations to test whether adaptive plasticity constrained subsequent genomic and phenotypic changes. Next, we evaluate how adaptive plasticity influences how well populations produced by each type of founder can evolve and retain novel adaptive traits. Finally, we examine lineages to determine whether adaptive plasticity facilitated the accumulation of cryptic genetic variation that would prove deleterious when the environment changed.

**Figure 2.**
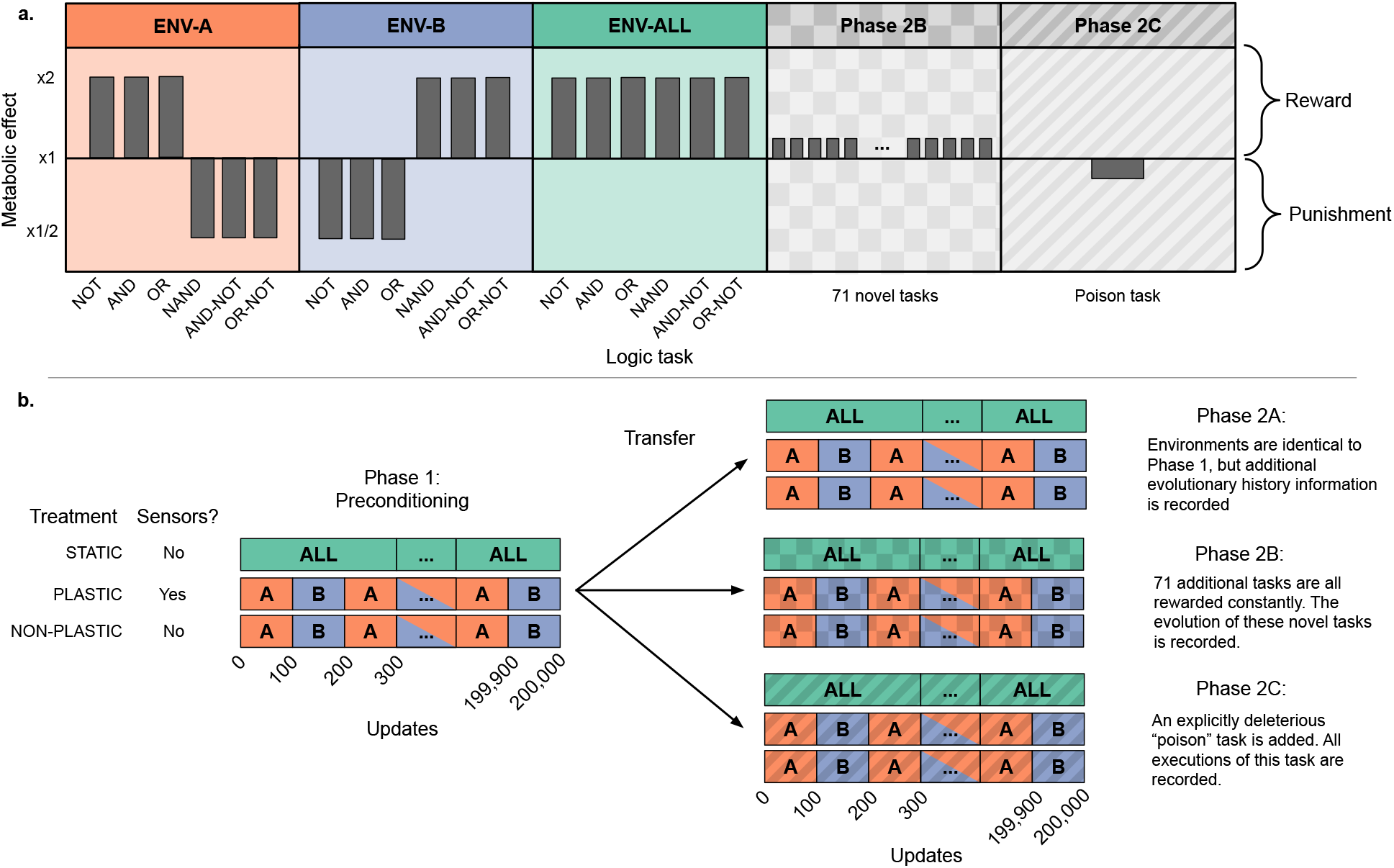
Overview of experimental design. The first three plots in panel (a) show the environments used in every experiment and whether they reward or punish each base task. Additionally, the last two subplots in (a) show the additional tasks added in phases 2B and 2C. All novel tasks in phase 2B confer a 10% metabolic reward, while executing the poisonous task in phase 2C causes a 10% metabolic punishment (bars not drawn to scale). Panel (b) shows treatment differences and experimental phases. Treatments are listed on the left, with each treatment specifying its environmental configuration and whether sensors are functional. We conducted three independent two-phase experiments, each described on the right. Phases 2B and 2C are textured to match their task definitions in panel (a). Phase one is repeated for *each* experiment with 100 replicate populations per treatment per experiment. For each replicate at the end of phase one, we used an organism of the most abundant genotype to found the second phase population. All STATIC and NON-PLASTIC populations move on to phase two, but PLASTIC populations only continue to the second phase if their most abundant genotype exhibits optimal plasticity. Metrics are recorded only in phase two.

We found that the evolution of adaptive plasticity reduced subsequent rates of evolutionary change in a cyclic environment. The non-plastic populations underwent more frequent selective sweeps and accumulated many more genetic changes over time, as non-plastic populations relied on genetic variation from *de novo* mutations to continuously readapt to environmental changes. The evolution of adaptive phenotypic plasticity buffered populations against environmental fluctuations, whereas repeated selective sweeps in non-plastic populations drove the accumulation of deleterious mutations and the loss of secondary beneficial traits via deleterious hitchhiking. As such, adaptively plastic populations were better able to retain novel traits than their non-plastic counterparts. In general, the evolution of adaptive phenotypic plasticity shifted evolutionary dynamics to be more similar to that of populations evolving in a static environment than to non-plastic populations evolving in an identical fluctuating environment.

## 2 Materials and Methods

### 2.1 The Avida Digital Evolution Platform

Avida is a study system wherein self-replicating computer programs (digital organisms) compete for space on a finite toroidal grid (Ofria et al., 2009). Each digital organism is defined by a linear sequence of program instructions (its genome) and a set of virtual hardware components used to interpret and express those instructions. Genomes are expressed sequentially except when the execution of one instruction (*e.g*., a “jump” instruction) deterministically changes which instruction should be executed next. Genomes are built using an instruction set that is both robust (*i.e*., any ordering of instructions is syntactically valid, though not necessarily meaningful) and Turing Complete (*i.e*., able to represent any computable function, though not necessarily in an efficient manner). The instruction set includes operations for basic computations, flow control (*e.g*., conditional logic and looping), input, output, and self-replication.

Organisms in Avida reproduce asexually by copying their genome instruction-by-instruction and then dividing. However, copy operations are imperfect and can result in single-instruction substitution mutations in an offspring’s genome. For this work, we configured copy operations to err at a rate of one expected mutation for every 400 instructions copied (*i.e*., a per-instruction error rate of 0.0025). We held individual genomes at a fixed length of 100 instructions; that is, we did not include insertion and deletion mutations. We used fixed-length genomes to control for treatment-specific conditions resulting in the evolution of substantially different genome sizes (Lalejini and Ferguson, 2021a)^1^, which could, on its own, drive differences in evolutionary outcomes among experimental treatments. When an organism divides in Avida, its offspring is placed in a random location on the toroidal grid, replacing any previous occupant. For this work, we used the default 60 by 60 grid size, which limits the maximum population size to 3600 organisms. As such, improvements to the speed of self-replication are advantageous in the competition for space.

During evolution, organism replication rates improve in two ways: by improving genome efficiency (*e.g*., using a more compact encoding) or by accelerating the rate at which the genome is expressed (their “metabolic rate”). An organism’s metabolic rate determines the speed at which it executes instructions in its genome. Initially, an organism’s metabolic rate is proportional to the length of its genome, but that rate is adjusted as it completes designated tasks, such as performing Boolean logic computations (Ofria et al., 2009). In this way, we can reward or punish particular phenotypic traits.

#### 2.1.1 Phenotypic plasticity in Avida

In this work, we measure a digital organism’s phenotype as the set of Boolean logic functions that it performs in a given environment. Sensory instructions in the Avida instruction set allow organisms to detect how performing a particular logic function would affect their metabolic rate (see supplemental material for more details, Lalejini and Ferguson 2021a). We define a phenotypically plastic organism as one that uses sensory information to alter which logic functions it performs based on the environment.

Phenotypic plasticity in Avida can be adaptive or non-adaptive for a given set of environments. Adaptive plasticity shifts net task expression closer to the optimum for the given environments. Non-adaptive plasticity changes task expression in either a neutral or deleterious way. In this work, optimal plasticity toggles tasks to always perfectly match the set of rewarded tasks for the given set of environments.

### 2.2 Experimental design

We conducted three independent experiments using Avida to investigate how the evolution of adaptive plasticity influences evolutionary outcomes in fluctuating environments. For each experiment, we compared the evolutionary outcomes of populations evolved under three treatments (Figure 2): (1) a **PLASTIC** treatment where the environment fluctuates, and digital organisms can use sensory instructions to differentiate between environmental states; (2) a **NON-PLASTIC** treatment with identical environment fluctuations, but where sensory instructions are disabled; and (3) a **STATIC** control where organisms evolve in a constant environment.

Each experiment was divided into two phases that each lasted for 200,000 updates^2^ of evolution (Figure 2), which is equivalent to approximately 30,000 to 40,000 generations. In phase one of each experiment, we preconditioned populations to their treatment-specific conditions. In phase two, we founded new populations with the evolved organisms from phase one and examined their subsequent evolution under given combinations of treatment and experimental conditions. During phase two, we tracked and saved each population’s evolutionary history as well as saving the full final population. Phase one was for preconditioning only; all comparisons between treatments were performed on phase two data.

#### 2.2.1 Environments

We constructed three experimental environments, abbreviated hereafter as “ENV-A”, “ENV-B”, and “ENV-ALL”. Figure 2 describes these environments based on whether each of six Boolean logic tasks (NOT, NAND, AND, OR-NOT, OR, and AND-NOT) is rewarded or punished. A rewarded task performed by an organism doubles their metabolic rate, allowing them to execute twice as many instructions in the same amount of time. A punished task halves an organism’s metabolic rate.

In both the PLASTIC and NON-PLASTIC treatments, the environment cycles between equal-length periods of ENV-A and ENV-B. Each of these periods persist for 100 updates (approximately 15 to 20 generations). Thus, populations experience a total of 1,000 full periods of ENV-A interlaced with 1,000 full periods of ENV-B during each experimental phase.

Organisms in the PLASTIC treatments differentiate between ENV-A and ENV-B by executing one of six sensory instructions, each associated with a particular logical task; these sensory instructions detect whether their associated task is currently rewarded or punished. By using sensory information in combination with execution flow-control instructions, organisms can conditionally perform different logic tasks depending on the current environmental conditions.

#### 2.2.2 Experiment Phase 1 – Environment preconditioning

For each treatment, we founded 100 independent populations from a common ancestral strain capable only of self-replication. At the end of phase one, we identified the most abundant (*i.e*., dominant) genotype and sampled an organism with that genotype from each replicate population to found a new population for phase two.

For the PLASTIC treatment, we measure plasticity by independently testing a given genotype in each of ENV-A and ENV-B. We discard phase one populations if the dominant genotype does not exhibit optimal plasticity. This approach ensures that measurements taken on PLASTIC-treatment populations during the second phase of each experiment reflect the evolutionary consequences of adaptive plasticity.

#### 2.2.3 Experiment Phase 2A – Evolutionary change rate

Phase 2A continued exactly as phase one, except we tracked the rates of evolutionary change in each of the PLASTIC-, NON-PLASTIC-, and STATIC-treatment populations. Specifically, we quantified evolutionary change rates using four metrics (each described in Table 1): (1) coalescence event count, (2) mutation count, (3) phenotypic volatility, and (4) mutational robustness.

**Table 1.**
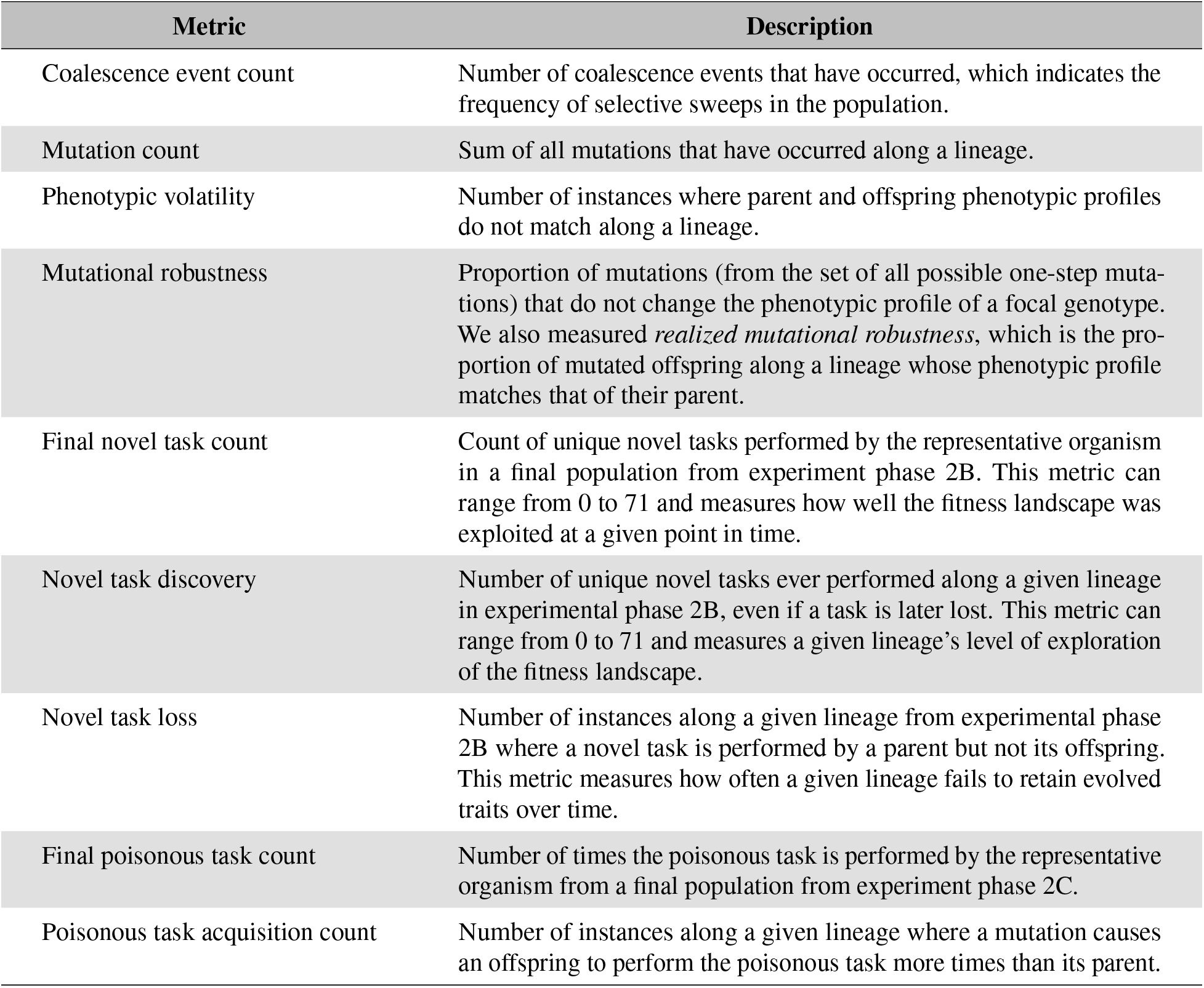
Metric descriptions.

| Metric | Description |
| --- | --- |
| Coalescence event count | Number of coalescence events that have occurred, which indicates the frequency of selective sweeps in the population. |
| Mutation count | Sum of all mutations that have occurred along a lineage. |
| Phenotypic volatility | Number of instances where parent and offspring phenotypic profiles do not match along a lineage. |
| Mutational robustness | Proportion of mutations (from the set of all possible one-step mutations) that do not change the phenotypic profile of a focal genotype. We also measured *realized mutational robustness*, which is the proportion of mutated offspring along a lineage whose phenotypic profile matches that of their parent. |
| Final novel task count | Count of unique novel tasks performed by the representative organism in a final population from experiment phase 2B. This metric can range from 0 to 71 and measures how well the fitness landscape was exploited at a given point in time. |
| Novel task discovery | Number of unique novel tasks ever performed along a given lineage in experimental phase 2B, even if a task is later lost. This metric can range from 0 to 71 and measures a given lineage’s level of exploration of the fitness landscape. |
| Novel task loss | Number of instances along a given lineage from experimental phase 2B where a novel task is performed by a parent but not its offspring. This metric measures how often a given lineage fails to retain evolved traits over time. |
| Final poisonous task count | Number of times the poisonous task is performed by the representative organism from a final population from experiment phase 2C. |
| Poisonous task acquisition count | Number of instances along a given lineage where a mutation causes an offspring to perform the poisonous task more times than its parent. |

#### 2.2.4 Experiment Phase 2B – Novel task evolution

Phase 2B extended the conditions of phase one by adding 71 novel Boolean logic tasks, which were always rewarded in all treatments (Ofria et al., 2009). The original six phase one tasks (NOT, NAND, AND, OR-NOT, OR, and AND-NOT; hereafter called “base” tasks) continued to be rewarded or punished according to the particular treatment conditions. An organism’s metabolic rate was increased by 10% for each novel task that it performed (limited to one reward per task). This reward provided a selective pressure to evolve these tasks, but their benefits did not overwhelm the 100% metabolic rate increase conferred by rewarded base tasks. As such, populations in the PLASTIC and NON-PLASTIC treatments could not easily escape environmental fluctuations by abandoning the fluctuating base tasks.

During this experiment, we tracked the extent to which populations evolving under each treatment were capable of acquiring and retaining novel tasks. Specifically, we used three metrics (each described in Table 1): (1) final novel task count, (2) novel task discovery, and (3) novel task loss.

#### 2.2.5 Experiment Phase 2C – Deleterious instruction accumulation

Phase 2C extended the instruction set of phase one with a `poison` instruction. When an organism executes a `poison` instruction, it performs a “poisonous” task, which reduces the organism’s metabolic rate (and thus reproductive success) but does not otherwise alter the organism’s function. We imposed a 10% penalty each time an organism performed the poisonous task, making the `poison` instruction explicitly deleterious to execute. We did not limit the number of times that an organism could perform the poisonous task, and as such, organisms could perform the poisonous task as many times as they executed the `poison` instruction.

We tracked the number of times each organism along the dominant lineage performed the poisonous task. Specifically, we used two metrics (each described in Table 1): (1) final poisonous task count and (2) poisonous task acquisition count.

### 2.3 Experimental analyses

For each of our experiments, we tracked and analyzed the phylogenetic histories of evolving populations during phase two. For each replicate, we identified an organism with the most abundant genotype in the final evolved population, and we used it as a *representative organism* for further analysis. We used the lineage from the founding organism to the representative organism as the *representative lineage* for further analysis. We manually inspected evolved phylogenies and found no evidence that any of our experimental treatments supported long-term coexistence. As such, each of the representative lineages reflect the majority of evolutionary history from a given population at the end of our experiment.

Some of our metrics (Table 1) required us to measure genotype-by-environment interactions. Importantly, in the fluctuating environments, we needed to differentiate phenotypic changes that were caused by mutations from those that were caused by environmental changes. To accomplish this, we produced organisms with the given focal genotype, measured their phenotype in each environment, and aggregated the resulting phenotypes to create a *phenotypic profile*. Although organisms with different genotypes may express the same set of tasks across environments, their phenotypic profiles may not necessarily be the same. For example, an organism that expresses NOT in ENV-A and NAND in ENV-B has a distinct phenotypic profile from one that expresses NAND in ENV-A and NOT in ENV-B.

While most analyses employed here are retrospective metrics applied to lineages, digital evolution allows precise manipulations on individual organisms and genomes. Mutational robustness uses this technique when looking at the possible mutations on a representative genotype. Genomes in Avida are linear sequences of instructions, and as such possible mutations can be simulated by substituting other instructions at the desired site. Indeed, the mutational robustness of a genotype examines all one-step mutations (*i.e*., each mutation where exactly one instruction is substituted). This allows us the disentangle whether results of the lineage metrics are a consequence of evolved genetic architectures or otherwise.

### 2.4 Statistical analyses

Across all of our experiments, we differentiated between sample distributions using non-parametric statistical tests. For each major analysis, we first performed a Kruskal-Wallis test (Kruskal and Wallis, 1952) to determine if there were significant differences in results from the PLASTIC, NON-PLASTIC, and STATIC treatments (significance level *α* = 0.05). If so, we applied a Wilcoxon rank-sum test (Wilcoxon, 1992) to distinguish between pairs of treatments. We applied Bonferroni corrections for multiple comparisons (Rice, 1989) where appropriate.

### 2.5 Software availability

We conducted our experiments using a modified version of the Avida software, which is open source and freely available on GitHub (Lalejini and Ferguson, 2021a). We used Python for data processing, and we conducted all statistical analyses using R version 4 (R Core Team, 2021). We used the tidyverse collection of R packages (Wickham et al., 2019) to wrangle data, and we used the following R packages for analysis, graphing, and visualization: ggplot2 (Wickham et al., 2020), cowplot (Wilke, 2020), Color Brewer (Harrower and Brewer, 2003; Neuwirth, 2014), rstatix (Kassambara, 2021), ggsignif (Ahlmann-Eltze and Patil, 2021), scales (Wickham and Seidel, 2020), Hmisc (Harrell Jr et al., 2020), fmsb (Nakazawa, 2019), and boot (Canty and Ripley, 2019). We used R markdown (Allaire et al., 2020) and bookdown (Xie, 2020) to generate web-enabled supplemental material. All of the source code for our experiments and analyses, including configuration files and guides for replication, can be found in our supplemental material, which is hosted on GitHub (Lalejini and Ferguson, 2021a). Additionally, our experimental data is available on the Open Science Framework at https://osf.io/sav2c/ (Lalejini and Ferguson, 2021b).

## 3 Results

### 3.1 Adaptive phenotypic plasticity slows evolutionary change in fluctuating environments

In experimental phase 2A, we tested whether adaptive phenotypic plasticity constrained or promoted subsequent evolutionary change in a fluctuating environment. First, we compared the total amount of evolutionary change in populations evolved under the PLASTIC, NON-PLASTIC, and STATIC treatments as measured by coalescence event count, mutation count, and phenotypic volatility (Figure 3). According to each of these metrics, NON-PLASTIC populations experienced a larger magnitude of evolutionary change than either PLASTIC or STATIC populations. We observed significantly higher coalescence event counts in NON-PLASTIC populations than in PLASTIC or STATIC populations (Figure 3a). NON-PLASTIC lineages had significantly higher mutation counts (Figure 3b) and phenotypic volatility than PLASTIC or STATIC lineages (Figure 3c).

**Figure 3.**
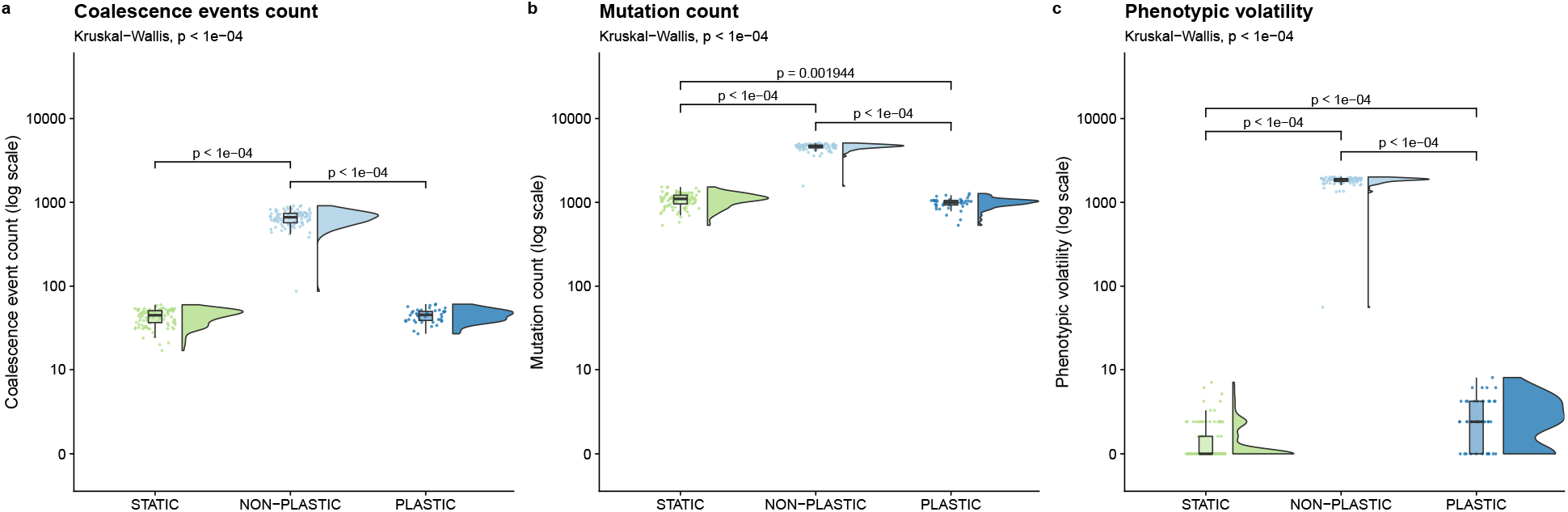
Magnitude of evolutionary change. Raincloud plots (Allen et al., 2019) of (a) coalescence event count, (b) mutation count, and (c) phenotypic volatility. See Table 1 for descriptions of each metric. Each plot is annotated with statistically significant comparisons (Bonferroni-corrected pairwise Wilcoxon rank-sum tests). Note that adaptive phenotypic plasticity evolved in 42 of 100 replicates from the PLASTIC treatment during phase one of this experiment; we used this more limited group to found 42 phase-two PLASTIC replicates from which we report these PLASTIC data.

Changing environments have been shown to increase generational turnover in Avida populations (Canino-Koning et al., 2016), which could explain why we observe a larger magnitude of evolutionary change at the end of 200,000 updates of evolution in NON-PLASTIC populations. Indeed, we found that significantly more generations of evolution elapsed in NON-PLASTIC populations (mean of 41090 ± 2702 std. dev.) than in PLASTIC (mean of 31016 ± 2615 std. dev.) or STATIC (mean of 30002 ± 3011 std. dev.) populations during phase 2A (corrected Wilcoxon rank-sum tests, p < 10^-4^).

To evaluate whether increased generational turnover explains the greater magnitude of evolutionary change in NON-PLASTIC populations, we examined the average number of generations between coalescence events and the realized mutational robustness of lineages (Table 1). A coalescence event indicates a selective sweep, which is a hallmark of adaptive evolutionary change. Realized mutational robustness measures the frequency that mutations cause phenotypic changes along a lineage. We expect that static conditions should favor fit lineages with high realized mutational robustness that no longer undergo rapid adaptive change and hence do not trigger frequent coalescence events. Under fluctuating conditions, however, lineages must be composed of plastic organisms if they are to maintain both high fitness and realized mutational robustness. Without plasticity, we expect fluctuating conditions to produce lineages with low realized mutational robustness and frequent coalescence events as populations must continually acquire and fix mutations to readapt to the environment.

On average, significantly fewer generations elapsed between coalescence events in NON-PLASTIC populations than in either PLASTIC or STATIC populations (Figure 4a). We also found that both STATIC and PLASTIC lineages exhibited higher realized mutational robustness relative to that of NON-PLASTIC lineages (Figure 4b); that is, mutations observed along NON-PLASTIC lineages more often caused phenotypic changes in offspring. Overall, our results indicate that NON-PLASTIC populations underwent more rapid (and thus a greater amount of) evolutionary change than either PLASTIC or STATIC populations.

**Figure 4.**
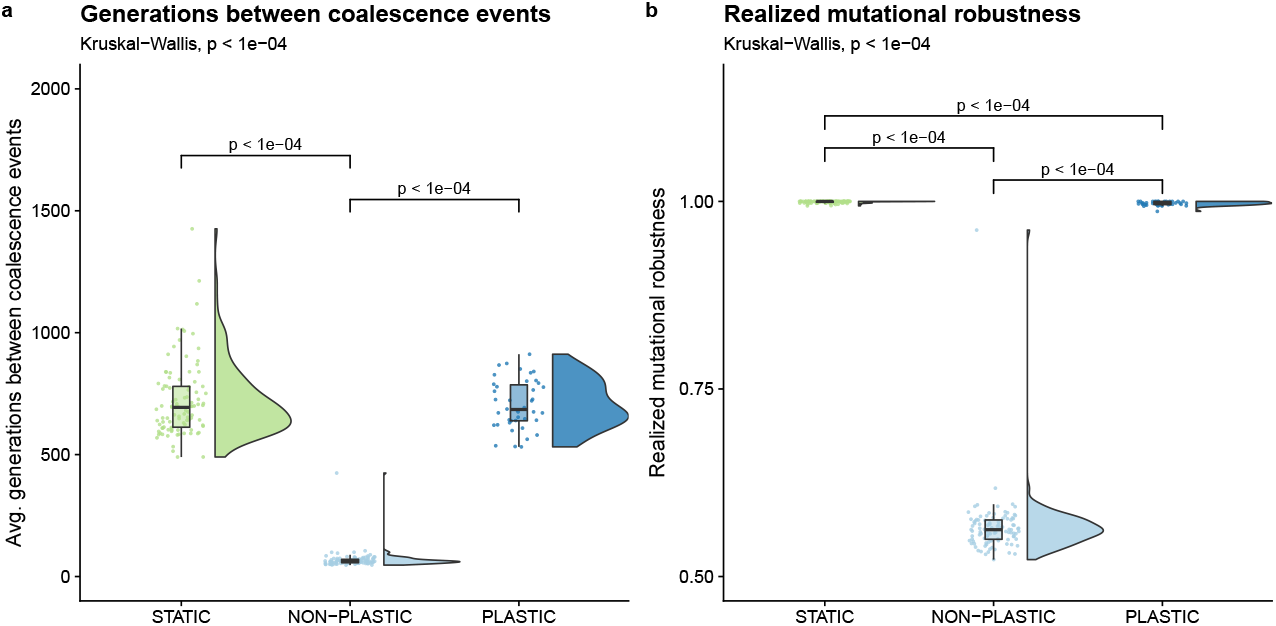
Pace of evolutionary change. Raincloud plots of (a) average number of generations between coalescence events, and (b) realized mutational robustness (Table 1). Each plot is annotated with statistically significant comparisons (Bonferroni-corrected pairwise Wilcoxon rank-sum tests).

While both STATIC and PLASTIC lineages exhibited high realized mutational robustness, we found that STATIC lineages exhibited higher realized robustness than PLASTIC lineages (Figure 4b). Overall, there were rare instances of mutations that caused a change in phenotypic profile across all PLASTIC lineages. Of these mutations, we found that over 80% (83 out of 102) of changes to phenotypic profiles were cryptic. That is, the mutations affected traits that would not have been expressed in the environment that the organism was born into but would have been expressed had the environment changed.

Given that NON-PLASTIC lineages exhibited the lowest realized mutational robustness of our three experimental treatments, we sought to determine if this effect was driven by differences in evolved genetic architectures. Specifically, did the NON-PLASTIC genetic architectures evolve such that mutations were more likely to result in phenotypic change? Such a mutational bias would trade off descendant fitness in the same environment in exchange for a chance of increasing descendant fitness in alternate environments. This strategy would be an example of diversifying bet-hedging (*i.e*., reducing expected mean fitness to lower variance in fitness) (Childs et al., 2010). Alternatively, the lower realized mutational robustness in NON-PLASTIC lineages could be due to survivorship bias, as we measured realized mutational robustness as the fraction of mutations observed along *successful* lineages that caused a phenotypic change.

We analyzed the mutational robustness of representative genotypes by calculating the fraction of single-instruction mutations that change the phenotypic profile. We found that mutations to representative genotypes on NON-PLASTIC lineages are *less* likely to result in a phenotypic change than mutations to comparable genotypes on either STATIC or PLASTIC lineages (Figure 5). These data provide evidence against NON-PLASTIC lineages engaging in a mutation-driven bet-hedging strategy, and instead, are consistent with the hypothesis that lower realized mutational robustness in the NON-PLASTIC treatment was due to survivorship bias.

**Figure 5.**
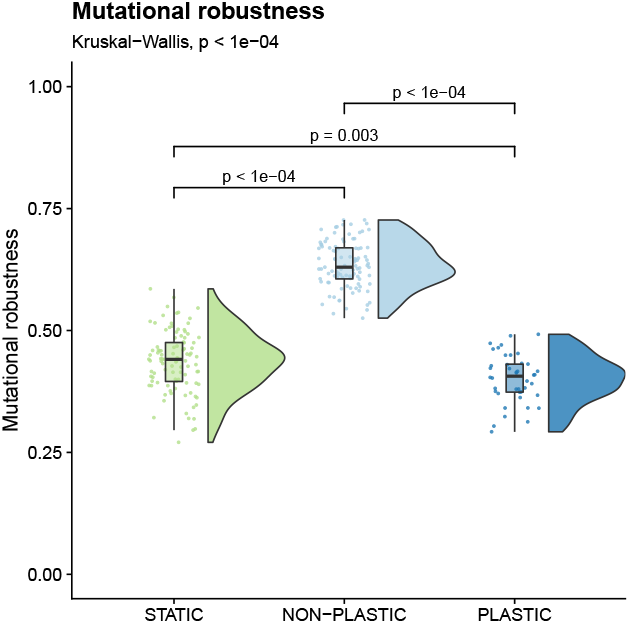
Mutational robustness. Raincloud plot of mutational robustness of each representative genotype (Table 1). The plot is annotated with statistically significant comparisons (Bonferroni-corrected pairwise Wilcoxon rank-sum tests).

In general, adaptive plasticity stabilized PLASTIC-treatment populations against environmental fluctuations, and their evolutionary dynamics more closely resembled those of populations evolving in a static environment. We observed no significant difference in the number and frequency of coalescence events in PLASTIC and STATIC populations. We did, however, observe small, but statistically significant, differences in each of the following metrics: elapsed generations, mutation counts, mutational robustness, and realized mutational robustness between PLASTIC and STATIC populations.

### 3.2 Adaptively plastic populations retain more novel tasks than non-plastic populations in fluctuating environments

We have so far shown that adaptive plasticity constrains the rate of evolutionary change in fluctuating environments. However, it is unclear how this dynamic influences the evolution of novel tasks. Based on their relative rates of evolutionary change, we might expect NON-PLASTIC-treatment populations to evolve more novel tasks than PLASTIC-treatment populations. But, how much of the evolutionary change in NON-PLASTIC populations is useful for exploring novel regions of the fitness landscape versus continually rediscovering the same regions?

To answer this question, we quantified the number of novel tasks performed by a representative organism in the final population of each replicate. We found that both PLASTIC and STATIC populations had significantly higher final task counts than NON-PLASTIC populations at the end of the experiment (Figure 6a). The final novel task count in PLASTIC and STATIC lineages could be higher than that of the NON-PLASTIC lineages for several non-mutually exclusive reasons. One possibility is that PLASTIC and STATIC lineages could be exploring a larger area of the fitness landscape when compared to NON-PLASTIC lineages. Another possibility is that the propensity of the NON-PLASTIC lineages to maintain novel traits could be significantly lower than PLASTIC or STATIC lineages. When we looked at the total sum of novel tasks discovered by each of the PLASTIC, STATIC, and NON-PLASTIC lineages, we found that NON-PLASTIC lineages generally explored a larger area of the fitness landscape (Figure 6b). Although the NON-PLASTIC lineages discovered more novel tasks, those lineages also exhibited significantly higher novel task loss when compared to PLASTIC and STATIC lineages (Figure 6c).

**Figure 6.**
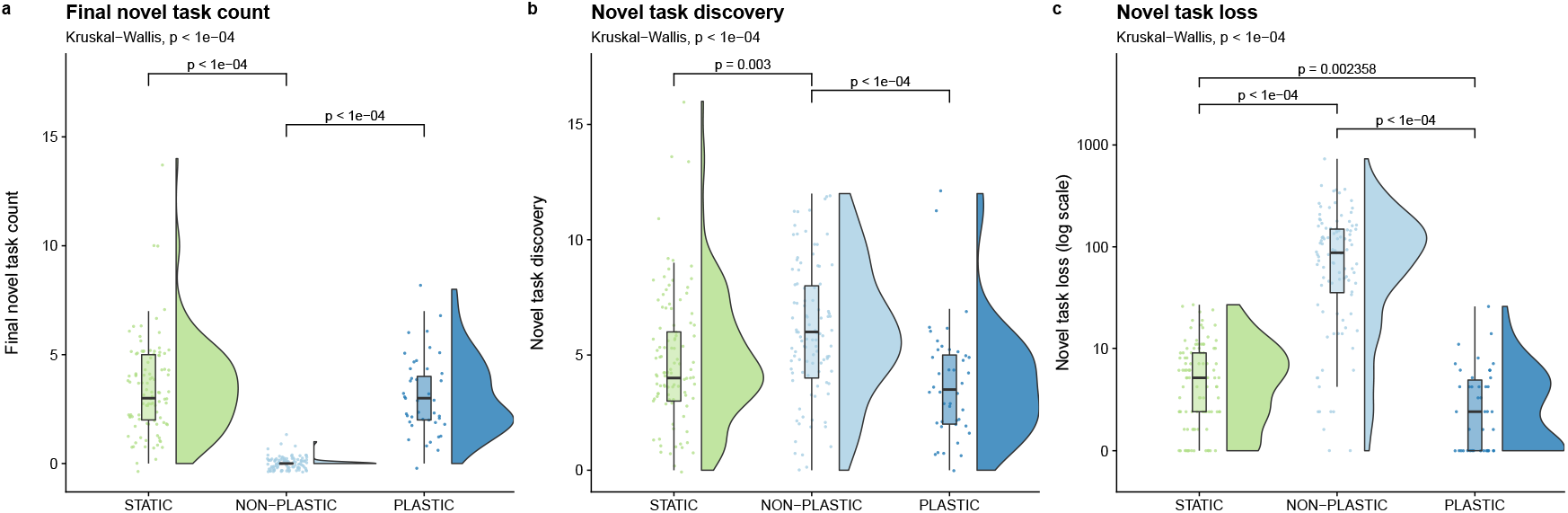
Novel task evolution. Raincloud plots of (a) final novel task count, (b) novel task discovery, and (c) novel task loss. See Table 1 for descriptions of each metric. Each plot is annotated with statistically significant comparisons (Bonferroni-corrected pairwise Wilcoxon rank-sum tests). Note that adaptive phenotypic plasticity evolved in 42 of 100 replicates from the PLASTIC treatment during phase one of this experiment; we used this more limited group to seed the resulting 42 phase-two PLASTIC replicates.

A larger number of generations elapsed in NON-PLASTIC populations than in PLASTIC or STATIC populations during our experiment (Lalejini and Ferguson, 2021a). Are NON-PLASTIC lineages discovering and losing novel tasks more frequently than PLASTIC or STATIC lineages, or are our observations a result of differences in generational turnover? To answer this question, we converted the metrics of novel task discovery and novel task loss to rates by dividing each metric by the number of elapsed generations along the associated representative lineages. We found no significant difference in the frequency of novel task discovery between NON-PLASTIC and STATIC lineages, and we found that PLASTIC lineages had a lower frequency of novel task discovery than STATIC lineages (Figure 7a). Therefore, we cannot reject the possibility that the larger magnitude of task discovery in NON-PLASTIC lineages was driven by a larger number of elapsed generations. NON-PLASTIC lineages had a higher frequency of task loss than either PLASTIC or STATIC lineages, and PLASTIC lineages tended to have a lower frequency of novel task loss than STATIC lineages (Figure 7b).

**Figure 7.**
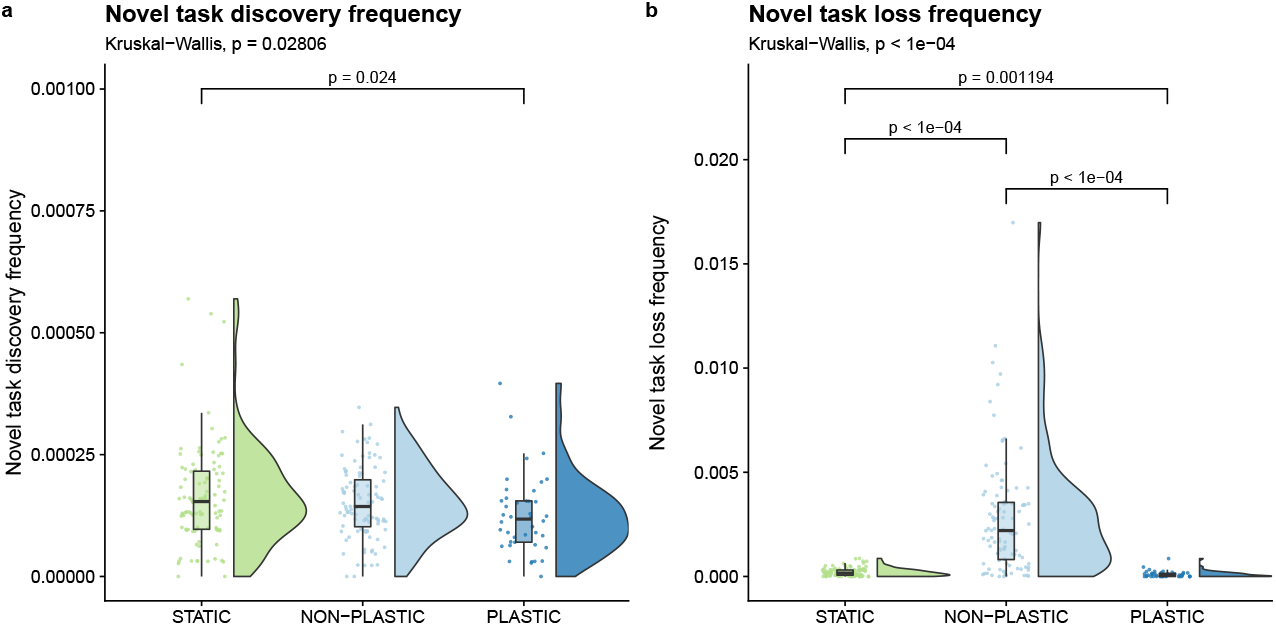
Rate of novel task evolution. Raincloud plots of (a) novel task discovery frequency and (b) novel task loss frequency. Each plot is annotated with statistically significant comparisons (Bonferroni-corrected pairwise Wilcoxon rank-sum tests).

Next, we examined the frequency at which novel task loss along lineages co-occurred with the loss or gain of any of the six base tasks. Across all NON-PLASTIC representative lineages, over 97% (10998 out of 11229) of instances of novel task loss co-occurred with a simultaneous change in base task profile. In contrast, across all PLASTIC and STATIC dominant lineages, we observed that approximately 20% (29 out of 142) and 2% (13 out of 631), respectively, of instances of novel task loss co-occurred with a simultaneous change in base task profile. As such, the losses of novel tasks in NON-PLASTIC lineages appear to be primarily due to hitchhiking.

### 3.3 Lineages without plasticity that evolve in fluctuating environments express more deleterious tasks

Phenotypic plasticity allows for genetic variation to accumulate in genomic regions that are unexpressed, which could lead to the fixation of deleterious instructions in PLASTIC populations. However, in NON-PLASTIC lineages, we observe a higher rate of novel task loss, indicating that they may be more susceptible to deleterious mutations (Figure 7b).

Therefore, in experiment phase 2C, we tested whether adaptive phenotypic plasticity can increase the incidence of deleterious task performance. Specifically, we added an instruction that triggered a “poisonous” task and measured the number of times it was executed. Each execution of the `poison` instruction reduces an organism’s fitness by 10%. At the beginning of phase 2C, the `poison` instruction is not present in the population, as it was not part of the instruction set during phase one of evolution. Accordingly, if a `poison` instruction fixes in a population, it must be the result of evolutionary dynamics during phase 2C, including cryptic variation or hitchhiking.

At the end of our experiment, no representative organisms from the PLASTIC or STATIC treatments performed the poisonous task under any environmental condition; however, representative organisms in 14% of replicates of the NON-PLASTIC treatment performed the poisonous task at least once. NON-PLASTIC lineages contained significantly more mutations that conferred the poisonous task as compared to PLASTIC or STATIC lineages (Figure 8a), and these mutations occurred at a significantly higher frequency in NON-PLASTIC lineages (Figure 8b).

**Figure 8.**
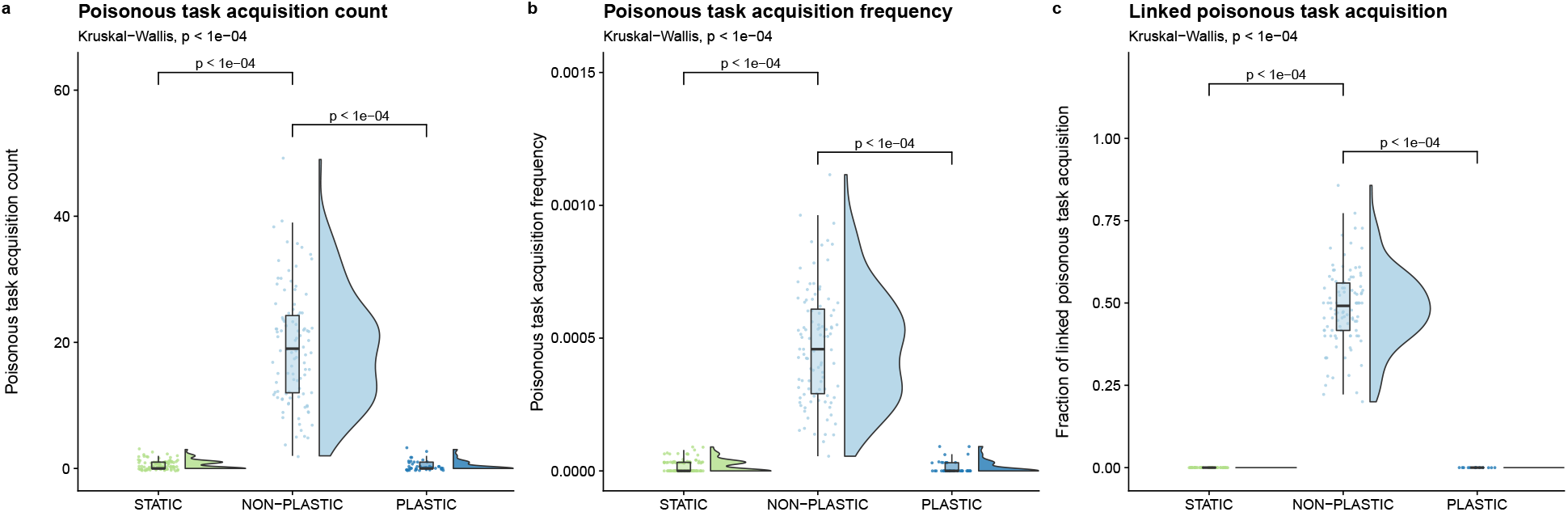
Deleterious instruction accumulation. Raincloud plots of (a) poisonous task acquisition, (b) poisonous task acquisition frequency, and (c) the proportion of mutations that increase poisonous task performance along a lineage that co-occur with a change in phenotypic profile. Each plot is annotated with statistically significant comparisons (Bonferroni-corrected pairwise Wilcoxon rank-sum tests). Note that adaptive phenotypic plasticity evolved in 43 of 100 replicates from the PLASTIC treatment during phase one of this experiment; we used this more limited group to seed the 43 phase-two PLASTIC replicates.

Next, we measured how often mutations that increased poisonous task performance co-occurred with changes to the base task profile within representative lineages. A poisonous instruction can fix in a lineage by having a beneficial effect that outweighs its inherent cost (*e.g*., knocking out a punished task) or through linkage with a secondary beneficial mutation at another site within the genome. Across all NON-PLASTIC representative lineages, we found that approximately 49% (956 out of 1916) of mutations that increased poisonous task expression co-occurred with a change in the base task profile (Figure 8c). In all representative lineages from the PLASTIC treatment, only 18 mutations increased poisonous task expression, and none co-occurred with a change in base task profile (Figure 8c). Likewise, only 58 mutations increased poisonous task performance in all representative lineages from the STATIC treatment, and none co-occurred with a change in base task profile (Figure 8c). We did not find compelling evidence that the few mutations that increased poisonous task expression occurred as cryptic variation in PLASTIC lineages.

We repeated this experiment with 3% and 30% metabolic rate penalties associated with the poisonous task, which produced results that were consistent with those reported here (Lalejini and Ferguson, 2021a).

## 4 Discussion

In this work, we used evolving populations of digital organisms to determine how adaptive phenotypic plasticity alters subsequent evolutionary dynamics and influences evolutionary outcomes in fluctuating environments. Specifically, we compared lineages of adaptively plastic organisms in fluctuating environments to both non-plastic organisms in those same environments and other non-plastic organisms in static environments.

### 4.1 Evolutionary change

We found strong evidence that adaptive plasticity slows evolutionary change in fluctuating environments. Adaptively plastic populations experienced fewer coalescence events and fewer total genetic changes relative to non-plastic populations evolving under identical environmental conditions (Figure 3). Whereas non-plastic populations relied on *de novo* mutations to adapt to each environmental fluctuation, plastic populations leveraged sensory instructions to regulate task performance. Indeed, in fluctuating environments, selection pressures toggle after each environmental change. We hypothesize that in non-plastic populations such toggling would repeatedly drive the fixation of mutations that align an organism’s phenotypic profile to the new conditions. This hypothesis is supported by the increased frequency of coalescence events in these populations (Figure 4a) as well as increased rates of genetic and phenotypic changes observed along the lineages of non-plastic organisms.

Representative lineages in the non-plastic treatment experienced lower realized mutational robustness than plastic and static lineages (Figure 4b). We reasoned that this lower realized mutational robustness was due to non-plastic populations evolving a bet-hedging strategy where mutations are more likely to modify the phenotypic profile. However, when we switched from measuring the realized mutational robustness of representative lineages to measuring the mutational robustness of representative genotypes (*i.e*., what fraction of one-step mutants change the phenotypic profile), we observed that non-plastic genotypes exhibited the highest mutational robustness of all three treatments (Figure 5). This result runs contrary to both our expectations and the results of other fluctuating environment studies in Avida (Canino-Koning et al., 2019). Canino-Koning et al. (2019) found that mutational robustness is negatively correlated with the number of task-encoding sites in the genome. In our work, most plastic and static genotypes encode all six base tasks, while most non-plastic genotypes only encode tasks from one environment; this results in fewer task-encoding sites, which may increase mutational robustness in non-plastic genotypes (relative to plastic and static genotypes). Regardless of the cause, this higher mutational robustness in non-plastic organisms indicates that bet-hedging is not driving the low realized mutational robustness observed in non-plastic lineages. Thus, we expect the lower realized mutational robustness in non-plastic lineages to be driven by survivorship bias. Because non-plastic lineages must rely on mutations to adapt to environmental changes, phenotype-altering mutations are often highly advantageous, and their selection decreases the realized mutational robustness of the lineage.

To our knowledge, this study is the first in-depth empirical investigation into how the de novo evolution of adaptive plasticity shifts the course of subsequent evolution in a cyclic environment. The relative rates of evolutionary change that we observed in non-plastic populations, however, are consistent with results from previous digital evolution studies. For example, Dolson et al. (2020) showed that non-plastic populations that were evolved in cyclically changing environments exhibited higher phenotypic volatility and accumulated more mutations than that of populations evolved under static conditions. Furthermore, Lalejini and Ofria (2016) visually inspected the evolutionary histories of non-plastic organisms evolved in fluctuating environments, observing that mutations along successful lineages readily switched the set of traits expressed by offspring.

Our results are also consistent with conventional evolutionary theory. A trait’s evolutionary response to selection depends on the strength of directional selection and on the amount of genetic variation for selection to act upon (Lande and Arnold, 1983; Zimmer and Emlen, 2013). In our experiments, non-plastic populations repeatedly experienced strong directional selection to toggle which tasks were expressed after each environmental change. As such, retrospective analyses of successful lineages revealed rapid evolutionary responses (that is, high rates of genetic and phenotypic changes). Evolved adaptive plasticity shielded populations from strong directional selection when the environment changed by eliminating the need for a rapid evolutionary response to toggle task expression. Indeed, both theoretical and empirical studies have shown that adaptive plasticity can constrain evolutionary change by weakening directional selection on evolving populations (Price et al., 2003; Paenke et al., 2007; Ghalambor et al., 2015).

### 4.2 The evolution and maintenance of novel tasks

In fluctuating environments, non-plastic populations explored a larger area of the fitness landscape than adaptively plastic populations (Figure 6b). However, adaptively plastic populations better exploited the fitness landscape, retaining a greater number of novel tasks than non-plastic populations evolving under identical environmental conditions (Figure 6a). In our experiment, novel tasks were less important to survival than the fluctuating base tasks. In non-plastic populations, when a mutation changes a base task to better align with current environmental conditions, its benefit will often outweigh the cost of losing one or more novel tasks. Indeed, we found that along non-plastic representative lineages, 97% of the mutations associated with novel task loss co-occurred with phenotypic changes that helped offspring adapt to current environmental conditions.

Previous studies have shown that transitory environmental changes can improve overall fitness landscape exploration in evolving populations of non-plastic digital organisms (Nahum et al., 2017). Similarly, changing environments have been shown to increase the rate of evolutionary adaptation in simulated network models (Kashtan et al., 2007). In our system, however, we found that *repeated* fluctuations reduced the ability of non-plastic populations to maintain and exploit tasks; that said, we did find that repeated fluctuations may improve overall task discovery by increasing generational turnover. Consistent with our findings, Canino-Koning et al. (2019) found that non-plastic populations of digital organisms evolving in a cyclic environment maintained fewer novel traits than populations evolving in static environments.

Our results suggest that adaptive phenotypic plasticity can improve the potential for populations to exploit novel resources by stabilizing them against stressful environmental changes. The stability that we observe may also lend some support to the hypothesis that phenotypic plasticity can rescue populations from extinction under changing environmental conditions (Chevin et al., 2010).

Our data do not necessarily provide evidence for or against the genes as followers hypothesis. The genes as followers hypothesis focuses on contexts where plastic populations experience novel or abnormally stressful environmental change. However, in our system, environmental changes were cyclic (not novel), and no single environmental change was *abnormally* stressful. Further, the introduction of novel tasks during the second phase of the experiment merely added static opportunities for fitness improvement. This addition did not change the meaning of existing environmental cues, nor did it require those cues to be used in new ways.

### 4.3 The accumulation of deleterious alleles

We found that non-plastic lineages that evolved in a fluctuating environment exhibited both greater totals and higher rates of poisonous task acquisition than that of adaptively plastic lineages (Figure 8). In asexual populations without horizontal gene transfer, all co-occurring mutations are linked. As such, deleterious mutations linked with a stronger beneficial mutation (*i.e*., a driver) can sometimes “hitchhike” to fixation (Smith and Haigh, 1974; Van den Bergh et al., 2018; Buskirk et al., 2017). Natural selection normally prevents deleterious mutations from reaching high frequencies, as such mutants are outcompeted. However, when a beneficial mutation sweeps to fixation in a clonal population, it carries along any linked genetic material, including other beneficial, neutral, or deleterious mutations (Barton, 2000; Smith and Haigh, 1974). Therefore, we hypothesize that deleterious genetic hitchhiking drove `poison` instruction accumulation along non-plastic lineages in changing environments.

Across our experiments, the frequency of selective sweeps in non-plastic populations provided additional opportunities for genetic hitchhiking with each environmental change. Indeed, representative lineages from non-plastic populations in the cyclic environment exhibited higher mutation accumulation (Figure 3b), novel trait loss (Figure 6c), and poisonous task acquisition (Figure 8a) than their plastic counterparts. In aggregate, we found that many (~49%; 956 / 1916) mutations that increased `poison` instruction execution in offspring co-occurred with mutations that provided an even stronger benefit by adapting the offspring to an environmental change. We expect that an even larger fraction of these deleterious mutations were linked to beneficial mutations, but our analysis only counted mutations that co-occurred in the same generation.

Theory predicts that under relaxed selection deleterious mutations should accumulate as cryptic variation in unexpressed traits (Lahti et al., 2009). Contrary to this expectation, we did not find evidence of `poison` instructions accumulating as cryptic variation in adaptively plastic lineages. One possible explanation is that the period of time between environmental changes was too brief for variants carrying unexpressed `poison` instructions to drift to high frequencies before the environment changed, after which purifying selection would have removed such variants. Indeed, we would not expect drift to fix an unexpressed trait since we tuned the frequency of environmental fluctuations to prevent valuable traits from being randomly eliminated during the off environment. Additionally, plastic organisms in Avida usually adjust their phenotype by toggling the expression of a minimal number of key instructions, leaving little genomic space for cryptic variation to accumulate.

### 4.4 Limitations and future directions

Our work lays the groundwork for using digital evolution experiments to investigate the evolutionary consequences of phenotypic plasticity in a range of contexts. However, the data presented here are limited to the evolution of *adaptively* plastic populations. Future work might explore the evolutionary consequences of maladaptive and non-adaptive phenotypic plasticity (*e.g*., Leroi et al. 1994), which are known to bias evolutionary outcomes (Ghalambor et al., 2015). Additionally in our experiments, sensory instructions perfectly differentiated between ENV-A and ENV-B, and environmental fluctuations never exposed populations to entirely new conditions. These parameters have been shown to influence evolutionary outcomes (Li and Wilke, 2004; Boyer et al., 2021), which if relaxed in the context of further digital evolution experiments, may yield additional insights.

We focused our analyses on the lineages of organisms with the most abundant genotype in the final population. These successful lineages represented the majority of the evolutionary histories of populations at the end of our experiment, as populations did not exhibit long-term coexistence of different clades. Our analyses, therefore, gave us an accurate picture of what fixed in the population. We did not, however, examine the lineages of extinct clades. Future work will extend our analyses to include extinct lineages, giving us a more complete view of evolutionary history, which may allow us to better distinguish adaptively plastic populations from populations evolving in a static environment.

As with any wet-lab experiment, our results are in the context of a particular model organism: “Avidian” self-replicating computer programs. Digital organisms in Avida regulate responses to environmental cues using a combination of sensory instructions and conditional logic instructions (`if` statements). The `if` instructions conditionally execute a single instruction depending on previous computations and the state of memory. As such, plastic organisms in Avida typically regulate phenotypes by toggling the expression of a small number of key instructions as opposed to regulating cohorts of instructions under the control of a single regulatory sequence (Lalejini and Ferguson, 2021a). This bias may limit the accumulation of hidden genetic variation in Avida genomes. However, as there are many model biological organisms, there are many model digital organisms that have different regulatory mechanisms (*e.g.*, Lalejini and Ofria 2018) that should be used to test the generality of our results.

## Supplemental Material

The supplemental material for this article is hosted on GitHub and can be found online at https://github.com/amlalejini/evolutionary-consequences-of-plasticity (Lalejini and Ferguson, 2021a).

## Data Availability Statement

The datasets generated and analyzed for this study can be found on the Open Science Framework at https://osf.io/sav2c/ (Lalejini and Ferguson, 2021b).

## Author Contributions

AL and AJF designed the experiments, developed the necessary experiment software, conducted experiments, analyzed the results, and drafted the manuscript. AL, AJF, NAG and CO edited and approved the manuscript.

## Acknowledgments

We thank members of the MSU Digital Evolution Lab for helpful comments and suggestions on this work. We also thank Luis Zaman whose keynote talk at the 2020 Artificial Life conference inspired the initial exploratory experiments from which this work blossomed. MSU provided computational resources through the Institute for Cyber-Enabled Research. Any opinions, findings, and conclusions or recommendations expressed in this material are those of the author(s) and do not necessarily reflect the views of the National Science Foundation, Department of Energy, Michigan State University, or The University of Idaho.

## Footnotes

1 We repeated our experiments without genome size restrictions and observed qualitatively similar results (see supplemental material, Lalejini and Ferguson 2021a).

2 One update in Avida is the amount of time required for the average organism to execute 30 instructions. See (Ofria et al., 2009) for more details.

